# OMCD: OncoMir Cancer Database

**DOI:** 10.1101/279125

**Authors:** Aaron Sarver, Anne Sarver, Ce Yuan, Subbaya Subramanian

## Abstract

MicroRNAs (miRNAs) are crucially important in the development of cancer. Their dysregulation, commonly observed in various types of cancer, is largely cancer-dependent. Thus, to understand the tumor biology and to develop accurate and sensitive biomarkers, we need to understand pan-cancer miRNA expression. At the University of Minnesota, we developed the OncoMir Cancer Database (OMCD), hosted on a web server, which allows easy and systematic comparative genomic analyses of miRNA sequencing data derived from more than 9,500 cancer patients tissue samples available in the Cancer Genome Atlas (TCGA). OMCD includes associated clinical information and is searchable by organ-specific terms common to the TCGA. Freely available to all users (www.oncomir.umn.edu/omcd/), OMCD enables (1) simple visualization of TCGA miRNA sequencing data, (2) statistical analysis of differentially expressed miRNAs for each cancer type, and (3) exploration of miRNA clusters across cancer types.

**Database URL:** www.oncomir.umn.edu/omcd

## Background

MicroRNAs are small non-coding RNAs that regulate gene expression through posttranscriptional modifications, specifically by binding to the 3’ untranslated region (UTR) of the target messenger RNAs (1). Dysregulation of miRNAs has been associated with various types of cancer, such as colorectal cancer, lung cancer, lymphoma, glioblastoma, and osteosarcoma (2). Their largely cancer-dependent dysregulation makes miRNAs candidate biomarkers for diagnosis, classification, and prognosis, as well as potential therapeutic targets (2). Using miRNAs as biomarkers for diagnosis and classification has already been approved by the United States Food and Drug Administration (FDA) for lung, thyroid, and kidney cancers. They have also been approved by the FDA for identifying the primary site of other cancer types. However, because of the cancer dependent nature of miRNAs, we need to understand pan-cancer miRNA expression to have a comprehensive understanding of the tumor biology and to develop accurate and sensitive biomarkers.

The Cancer Genome Atlas (TCGA), a collaboration between the National Cancer Institute and the National Human Genome Research Institute, contains miRNA expression data for nearly 10,000 patients with 33 different cancer types (3). Currently, the 2 major web-based repositories of analyzed TCGA data are the cBioPortal and the Broad Institute’s genomic DNA affinity chromatography (GDAC) FireBrowse (4). However, both of those platforms focus mainly on the analysis and visualization of genomic and mRNA data; neither of them enables in-depth analysis or comparative visualization of miRNA data. Still another database, known as OncomiR, enables analysis of TCGA miRNA data and calculates miRNA markers as survival signatures (5). But OncomiR does not provide simple visualization of TCGA miRNA expression data nor the ability to explore miRNA clusters.

At the University of Minnesota, we developed the OncoMir Cancer Database (OMCD), which enables (1) simple visualization of TCGA miRNA sequencing data, (2) statistical analysis of differentially expressed miRNAs for each cancer type, and (3) exploration of miRNA clusters across cancer types.

## Construction and content

To create OMCD, we used the LAMP software bundle (Linux, Apache 2, MySQL 5.0, and PHP) and hypertext markup language (HTML), as described previously (6). Those applications are accessible to researchers across the globe. To host OMCD’s web application, we chose an Apache web server. To generate the user interface and enable communication with the MySQL database at the back end, we chose PHP, given its database-driven architecture that was designed for incorporation of additional information. Normalized expression data, statistical results, and annotation data are all stored in OMCD. To facilitate data retrieval and selection of different criteria for analysis, we designed a user-friendly graphic interface.

To construct the content of OMCD, we downloaded from TCGA the miRNA expression data of 9,656 patients (represented by 8,993 tumor samples and 663 control samples of normal tissue with 33 different cancer types (https://gdc.nci.nih.gov; **Table 1**). We used a 2-group *t* test to determine which miRNAs were differentially expressed between (1) a cancer patient’s control and tumor samples, for a given cancer type, (2) a cancer patient’s control sample, as compared with all other patients’ available control samples, and (3) a cancer patient’s tumor sample, as compared with all other patients’ available tumor samples. It can be noted that each of our 3 analyses had a different statistical power, which accounted for the absence of any miRNA from a specific dataset.

**Table 1:**
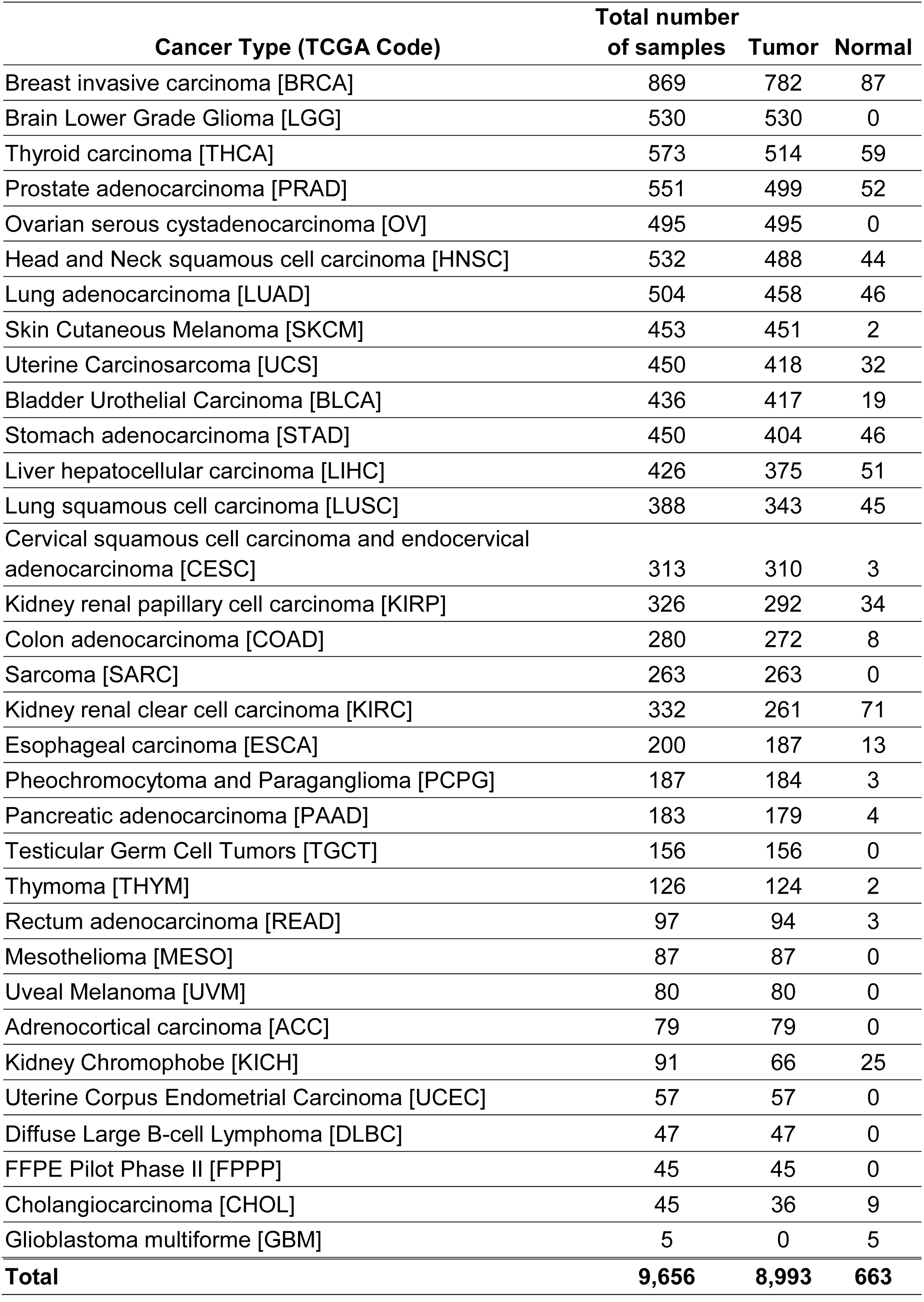
Number of patients in the OncoMir Cancer Database (OMCD), by cancer type

## Utility and discussion

Our newly developed OMCD is available at www.oncomir.umn.edu/omcd. It features 4 types of search function (**Figure 1A**). Here we use the miR-21 expression in colon adenocarcinoma (COAD) as an example. In the current version of OMCD repository, we have 8 control samples and 272 tumor samples for COAD. When we searched for miR-21 in COAD samples (**Figure 1A, B**), we obtained a heat map showing the absolute expression level of miR- 21 in all COAD samples (**Figure 1C**). We also obtained numeric expression data (**Figure 1D**; not completely shown, because of space limitations) and relative expression data (**Figure 1E**). When we clicked on “COAD”, we were taken to a page showing links to additional analysis (**Figure 1F**). Those links provided detailed information about the location of miR-21 and the names of colocalized miRNAs (miRNA clusters), as well as additional internal links to the expression data of miR-21 in other cancer types and to further statistical analysis (**Figure 1H**). The page also provides external links to the miRDB website for target prediction (www.mirdb.org) and to Google Scholar for literature searches (7). This page also enabled us to visualize colocalized miRNA expressions in a heat map showing absolute expression (**Figure 1G**). Expression levels of colocalized miRNAs can be displayed for all cancer types (not shown) and can be visualized in absolute and relative heat maps as well as in the form of numeric data. The 3 statistical analyses that we performed—using a cancer patient’s control vs. tumor samples; a cancer patient’s control sample vs. all other patients’ control samples; and a cancer patient’s tumor sample vs. all other patients’ tumor samples—allowed us to further visualize the expression patterns of miR-21 across different cancer types (**Figure 1H**).

**Figure 1:**
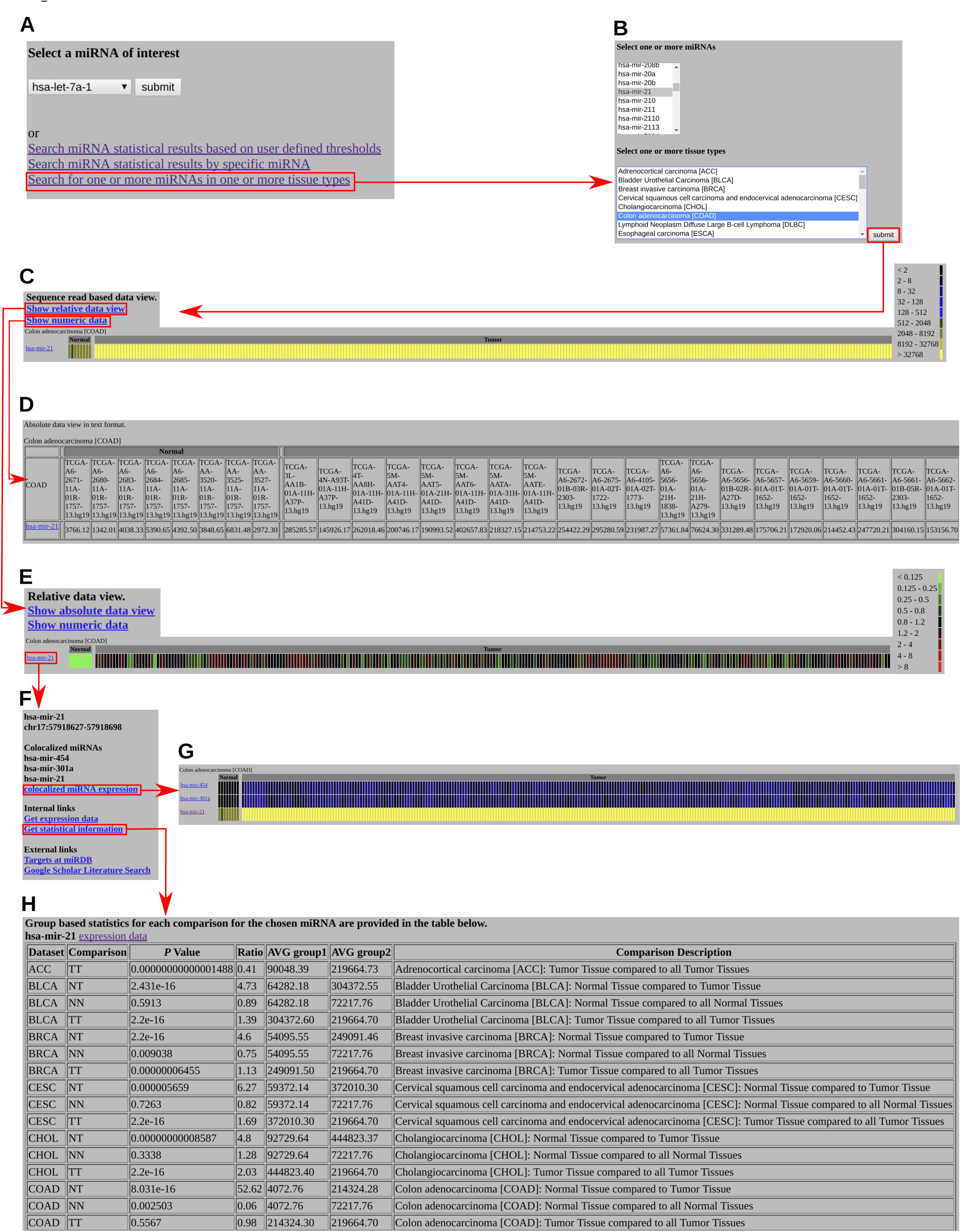
Screenshots of our sample analyses of miR-21 in COAD. **A, B.** Advanced Search options in OMCD, enabling searches by miRNA, by cancer types, and by statistical results. **C.** Heat map. **D.** Numeric view of absolute expressions of miR-21 in COAD control and tumor samples. **E.** Heat map of relative expression of miR-21 in COAD. **F.** Information and external links. **G.** Heat map of miR-21 cluster members, enabling exploration of the expression patterns of colocalized miRNAs. **H.** Statistical results of group-based comparisons of miR-21 in different cancer types.

To further demonstrate the utility of our database we identified miRNA which was recurrently significantly differentially expressed between tumor and normal control samples with a highly significant p-value < 0.000001 and an average fold change greater than the absolute value of 2 which were recurrently present in 5 or more tumor normal comparisons (**Figure 2**). Many miRNAs are well known in cancers and have been reported to be differentially expressed (between tumor and control samples) in a wide range of cancer types. For example, miR-21 is consistently upregulated in most cancer types (8). Thus, it could potentially be a *general* cancer biomarker, but it is not a suitable biomarker for *specific* cancer types.

**Figure 2:**
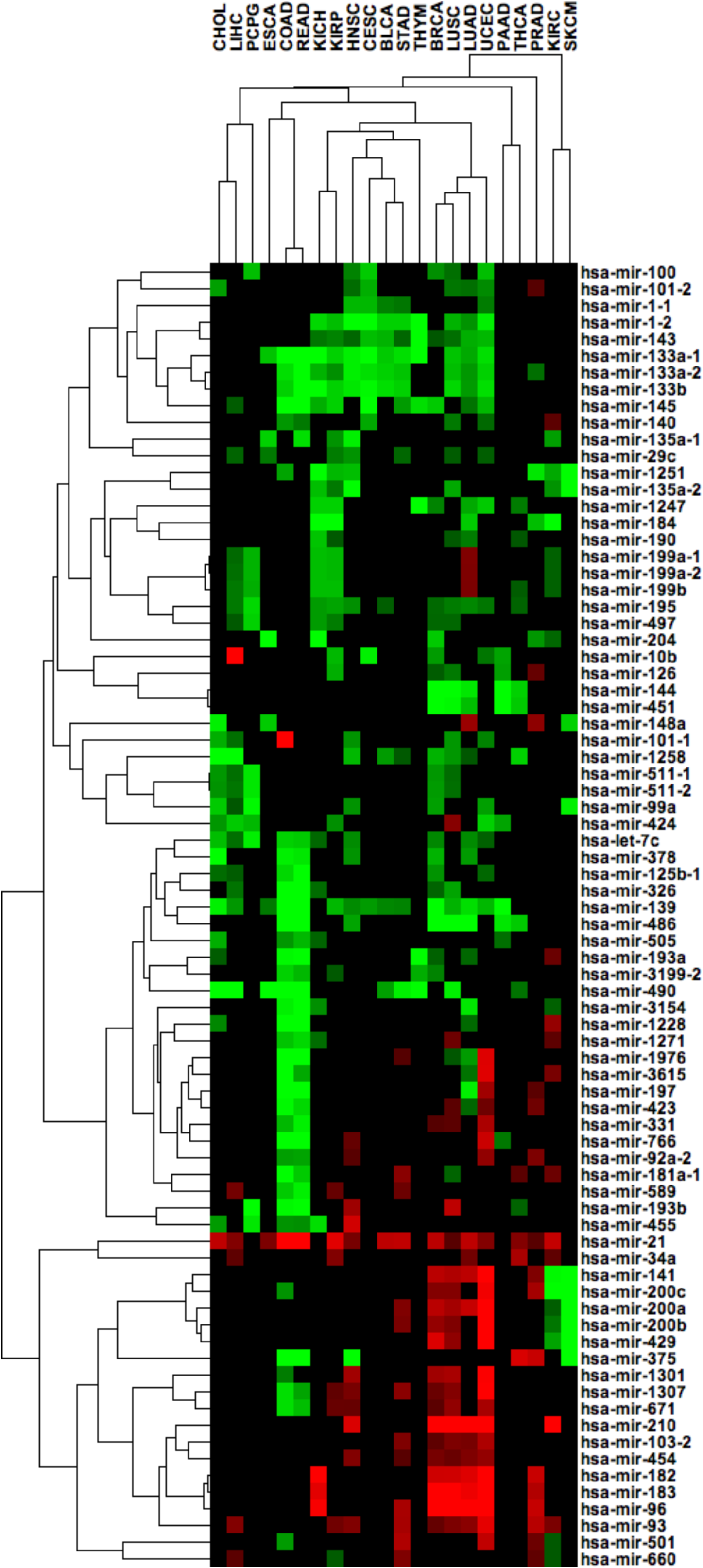
Heat map of differentially expressed miRNAs in tumor vs. control samples (*P* < 0.000001, with a mean fold change in the tumor samples greater than in 5 such comparisons). Red = upregulation; green = downregulation.

We also found that the COAD cluster and the rectal adenocarcinoma (READ) cluster had a very similar miRNA expression pattern, as compared with other cancer types. However, even though the COAD and READ clusters had a very similar miRNA expression pattern overall, miR-101-1 expression was significantly different between the COAD tumor and control samples—but *not* for the READ tumor and control samples **(Figure 2)**. Additionally, because miR-101 expression was not significantly higher in other cancer types, it is reasonable to hypothesize that this miRNA is a biomarker for COAD. Similarly, we found that miR-10b expression was significantly higher in hepatocellular carcinoma (LIHC), but not in other cancer types. These are just 2 examples of the potentially testable hypotheses that OMCD is able to generate. To more fully investigate the function of miR-101 in COAD and miR-10b in LIHC, further experimental validation is warranted.

## Discussion

Evidence from the past decade indicates that miRNAs play a crucial role in the development of various cancer types. With the advent of high-throughput sequencing technology, more high-throughput miRNA data are now publicly available. Our OMCD, which developed at the University of Minnesota, is a simple web-based repository that allows easy and systematic comparative genomic analyses of miRNA expression data.

In our OMCD testing, we were able to identify miR-101 as a biomarker candidate specifically for COAD. We found that its expression level was significantly higher in COAD tumors, but not in other tumors. Previous studies, however, showed miR-101 expression levels in colorectal cancer that were different from our results (9,10). Those previous studies suggested that miR-101 expression was downregulated in colorectal cancer and that it was a tumor-suppressing miRNA whose overexpression inhibited tumor invasion and growth (9,10). Interestingly, when we searched miR-101 expression in OncomiR (www.oncomir.org), which is also based on TCGA data, we again found that this miRNA was overexpressed in COAD tumors. Given the conflicting results for miR-101 in COAD in those 2 previous studies vs. our own use of both OMCD and OncomiR, further investigation into the function of miR-101 in COAD is needed, in order to definitively ascertain whether or not it is a suitable biomarker for COAD.

We also observed in our OMCD testing that miR-10b could be a potential biomarker for LIHC (11). Previous studies showed that miR-10b was indeed highly expressed in LIHC, that it was involved in neoplastic transformation of liver cancer stem cells, and promotes tumor metastasis (12–14). Other previous studies also showed an oncogenic role of miR-10b in breast cancer, gastric cancer, and glioblastoma (15–18). All of those studies suggest that miR-10b has a multifaceted function in many cancer types; further investigation is required, in order to definitively ascertain whether or not it is a suitable biomarker for LIHC.

Our current version of OMCD, derived from TCGA, contains the miRNA expression data of 9,656 patients (represented by 8,993 tumor samples and 663 control samples of normal tissue) with 33 different cancer types. To our knowledge, OncomiR (www.oncomir.org) is the only other TCGA-based online resource, besides OMCD, for analyzing miRNA expression data (5). A limitation of both OncomiR and our current version of OMCD is the lack of miRNA datasets from other cancer patient cohorts. But unlike OMCD, OncomiR lacks the option to analyze miRNA clusters. It is important to consider miRNA cluster members when studying miRNAs in cancers, especially to generate hypotheses from high-throughput data. Usually, miRNA cluster members have similar expression levels, but they potentially have vastly different biological functions. The ability to visualize and explore miRNA clusters in OMCD is crucial to develop defendable hypotheses.

In the future, we plan to expand OMCD by incorporating additional miRNA expression datasets from public data repositories such as Gene Expression Omnibus (GEO), Genomic Data Commons (GDC), and European Bioinformatics Institute (EBI). We believe this will significantly improve the ability to use OMCD to develop defendable hypotheses.

## List of Abbreviations

miRNAs – microRNAs

OMCD – OncoMir Cancer Database

TCGA – The Cancer Genome Atlas

FDA – United States Food and Drug Administration

GDAC – genomic DNA affinity chromatography

HTML – hypertext markup language

COAD – Colon adenocarcinoma

READ – Rectal adenocarcinoma

GEO – Gene Expression Omnibus

GDC – Genomic Data Commons

EBI – European Bioinformatics Institute

## Declarations

### Ethics approval and consent to participate

No ethics approval and content is required.

### Consent for publication

No consent for publication is required.

### Availability of data and material

The Cancer Genome Atlas data are available at gdc.nci.nih.gov. The OMCD database is freely accessable at www.oncomir.umn.edu/omcd.

### Funding

SS is supported by research grants funded by the National Cancer Institute of the National Institutes of Health, number R03CA219129; CY, by supported by Norman Wells Memorial Colorectal Cancer fellowship and Healthy Foods Healthy Lives Institute Graduate and Professional Student Research Grant of the University of Minnesota. The funding sources do not have any role in the design, data analysis, interpretation of data nor in writing the manusript. **Authors’ contributions**

Aaron S and SS, developed the concept; Aaron S and Anne S developed the database; CY, Aaron S and SS wrote the manuscript. All authors read and approved the final manuscript.

## Acknowledgments

We thank The Cancer Genome Atlas for providing unrestricted use of the microRNA data. We also thank Dr. Mary Knatterud for assisting in manuscript preparation.

